# Activity of Antibiotics against *Burkholderia cepacia* complex in Artificial Sputum Medium

**DOI:** 10.1101/2023.12.13.571502

**Authors:** Anusha Shukla, Shade Rodriguez, Thea Brennan-Krohn

## Abstract

*Burkholderia cepacia* complex (Bcc) is a collection of intrinsically drug-resistant gram-negative bacteria that cause life-threatening pulmonary disease in people with cystic fibrosis (CF). Standard antimicrobial susceptibility testing methods have poor predictive value for clinical outcomes in people with Bcc infections, likely due in part to the significant differences between *in vitro* testing conditions and the environment in which Bcc grow in the lungs of people with CF. We tested the activity of six commonly used antibiotics against two clinical Bcc strains grown to high density in an artificial sputum medium in order to assess their activity in conditions mimicking those found *in vivo*. There were major discrepancies between standard susceptibility results and activity in our model, with some antibiotics, including ceftazidime, showing minimal activity despite low MICs, while others, notably tobramycin, were more active in high-density growth conditions than in standard assays. This work underscores the urgent need to develop more clinically relevant susceptibility testing approaches for Bcc.

## INTRODUCTION

Cystic Fibrosis (CF) is a life-threatening multisystem disease primarily affecting the lungs, pancreas, and gastrointestinal tract (1) caused by autosomal recessive mutations in the gene encoding the cystic fibrosis transmembrane conductance regulator (CFTR) chloride channel protein. In the airways, the altered CFTR proteins cause decreased chloride secretion from epithelial cells and abnormal hydration of secretions (2). Thick mucus, and the resulting ciliary impairment, promote bacterial colonization. People with CF typically acquire bacterial colonization of the respiratory tract during childhood and require intermittent, prolonged courses of antibiotics to treat pulmonary exacerbations, which are characterized by acute worsening of respiratory symptoms and pulmonary function (3). The most common colonizing organisms are *Pseudomonas aeruginosa* and *Staphylococcus aureus*, but perhaps the most feared is the *Burkholderia cepacia* complex (Bcc) (4).

Bcc is a collection of gram-negative non-glucose fermenting bacteria initially described as consisting of nine phenotypically similar species, known as genomovars (5), but subsequently reclassified as comprising approximately 22 known species (6). Bcc is notorious for its propensity to spread rapidly amongst people with CF (7) and to cause fulminant, often fatal disease in the form of “cepacia syndrome” (8). Furthermore, Bcc infection in CF is associated with increased morbidity and mortality, and it is considered a contraindication for lung transplantation at many centers (7). Bcc also causes life-threatening infections in people with chronic granulomatous disease (9) and has caused healthcare-associated outbreaks in intensive care units (10, 11).

Treatment options for Bcc are limited due to its intrinsic resistance to many antibiotics, including aminoglycosides, polymyxins, and many beta-lactams. Isolates of Bcc also rapidly acquire additional resistance mechanisms once exposed to antibiotics (12). This shortage of intervention strategies is further compounded by the limitations of standard antimicrobial susceptibility testing (AST) methods for Bcc. The conditions under which AST is performed differ dramatically from the mucus layer in which bacteria grow in the lungs of people with CF, an environment which favors development of bacterial biofilms and aggregates (7). This profound discrepancy in growth conditions may explain the poor clinical predictive value of Bcc AST results (13, 14). Reproducibility of standard MIC testing in Bcc has also proved suboptimal (15). As a result of these concerns, the European Committee on Antimicrobial Susceptibility Testing (EUCAST) declines to promulgate interpretive criteria for Bcc (13). There is thus an urgent need to better understand which drugs are most likely to be effective in the CF lung and to determine which *in vitro* tests, if any, would be preferable to standard AST.

In order to evaluate the activity of antibiotics under conditions that better approximate those found in the lungs of people with CF, we assayed antibiotic activity against two clinical Bcc isolates grown in an artificial sputum medium characterized by increased concentrations of mucin, free DNA, and amino acids (16, 17). Growth in sputum or artificial sputum medium has been shown to affect gene expression and metabolite production of Bcc (18, 19), and we hypothesized that antibiotic activity would be affected by bacterial growth in this medium. We also allowed bacteria to grow to high density over 72 hours before adding antibiotic, in order to better simulate an established infection rather than the lower-density, exponential growth phase population typically used in standard AST (20) and time-kill studies.

## RESULTS

### MIC testing

Two clinical strains, *Burkholderia cenocepacia* K56-2 and *Burkholderia multivorans* CGD1, were evaluated; isolates from these species account for the majority of Bcc infections in people with CF (7). Modal MIC results for the two strains, based on 5 biological replicates, are shown in **Table 1**.

**Table 1.** Minimal Inhibitory Concentrations (MICs)

| Antibiotic | Modal MIC (µg/mL) |  |
| --- | --- | --- |
|  | *B. cenocepacia* K56-2 | *B. multivorans* CGD1 |
| Minocycline | 16 (R) | 8 (I) |
| Meropenem | 4 (S) | 16 (R) |
| Tobramycin | 32 (N/A) | 16 (N/A) |
| Ceftazidime | 2 (S) | 2 (S) |
| Levofloxacin | 4 (I) | 2 (S) |
| Trimethoprim-Sulfamethoxazole | 2/38 (S) | 1/19 (S) |

S: Susceptible; I: Intermediate; R: Resistant; N/A: Not available. Categories refer to Clinical and Laboratory Standards Institute (CLSI) interpretive criteria.

### Time-kill studies with established growth in ASMDM

Antibiotic activity was evaluated against bacteria growing in conditions designed to simulate the environment in the lungs of people with CF: bacteria were grown in ASMDM and were incubated for 72 hours to reach high-inoculum, stationary growth phase conditions before antibiotics were added (**Figs. 1, S1**).

**FIGURE 1.**
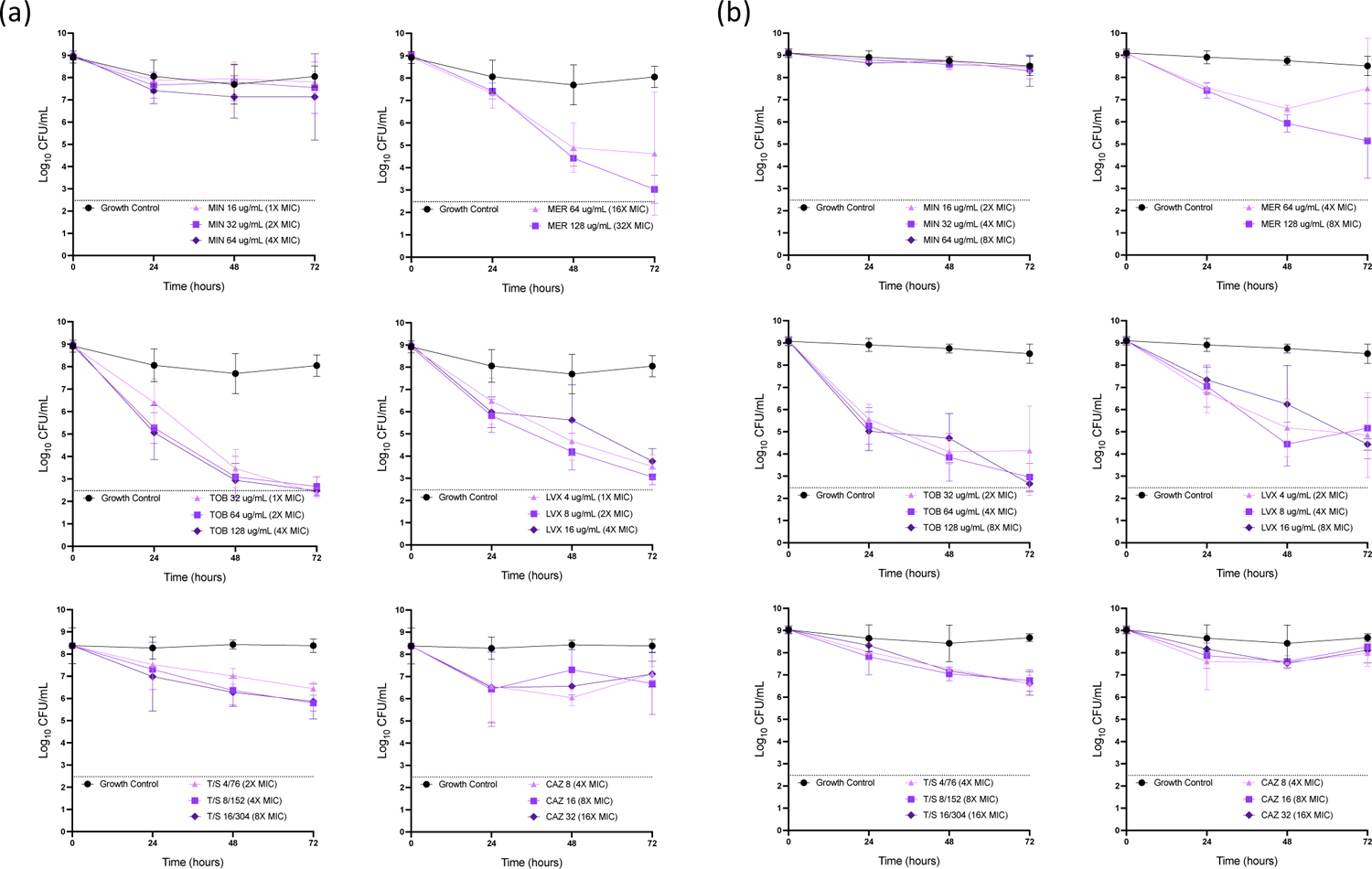
Time-kill studies in ASMDM with established growth of (a) *B. cenocepacia* K56-2 and (b) *B. multivorans* CGD1 MIN: minocycline; MER: meropenem; TOB: tobramycin; LVX: levofloxacin; T/S: trimethoprim-sulfamethoxazole; CAZ: ceftazidime. Graphs show geometric mean and standard deviation of ≥3 replicates. Dashed line indicates assay lower limit of detection. For assays performed simultaneously, the same growth control wells were used.

Activity in this environment was not predictable based on MIC results. For example, minocycline exhibited no killing activity against either strain even at concentrations up to 8X MIC. Similarly, ceftazidime caused less than a 1.9 log_10_ CFU/mL decline in colony count at any time point against either strain at 16X MIC. This finding was particularly notable as both strains were highly susceptible to ceftazidime by MIC testing. By contrast, at 2X MIC, tobramycin caused a decline in colony count of 6.3 log_10_ at 72 hours for *B. cenocepacia* and 5.0 log_10_ for *B. multivorans*, and levofloxacin reduced colony counts by 5.9 log_10_ at 72 hours for *B. cenocepacia* and 4.2 log_10_ for *B. multivorans*.

### Time-kill studies under different growth conditions

To evaluate the effect of different growth conditions and media types, a selected antibiotic concentration was tested against each strain grown in standard bacterial growth medium (CAMHB) for 72 hours before addition of antibiotics (“CAMHB Established growth”) and in each type of media under standard time-kill conditions, with antibiotics added at the beginning of the experiment (“ASMDM Standard time-kill” and “CAMHB Standard time-kill”) (**Figs. 2, 3**). In most cases antibiotic activity was similar between the two media types, but there were several cases in which antibiotic activity differed between established-growth conditions and standard time-kill conditions. For meropenem, killing was more pronounced in standard time-kill conditions, with a decline in colony count in ASMDM at 24 hours of 1.5-1.6 log_10_ CFU/mL in established growth assays but 2.8-3.8 log_10_ CFU/mL in standard time-kill assays. Trimethoprim-sulfamethoxazole also performed better against *B. multivorans* in standard time-kill assays. Somewhat unexpectedly, some drugs performed better in established growth assays. In standard time-kill studies in ASMDM, bacteria treated with tobramycin increased by >1 log_10_ CFU/mL by 72 hours, whereas in established growth studies, the colony count had fallen by >6 log_10_ CFU/mL at 72 hours. A similar pattern was seen with levofloxacin for *B. cenocepacia*, where growth at 72 hours had increased by 1.8 log_10_ CFU/mL in the standard-time kill study but fallen by 5.4 log_10_ CFU/mL in the established growth assay.

**FIGURE 2.**
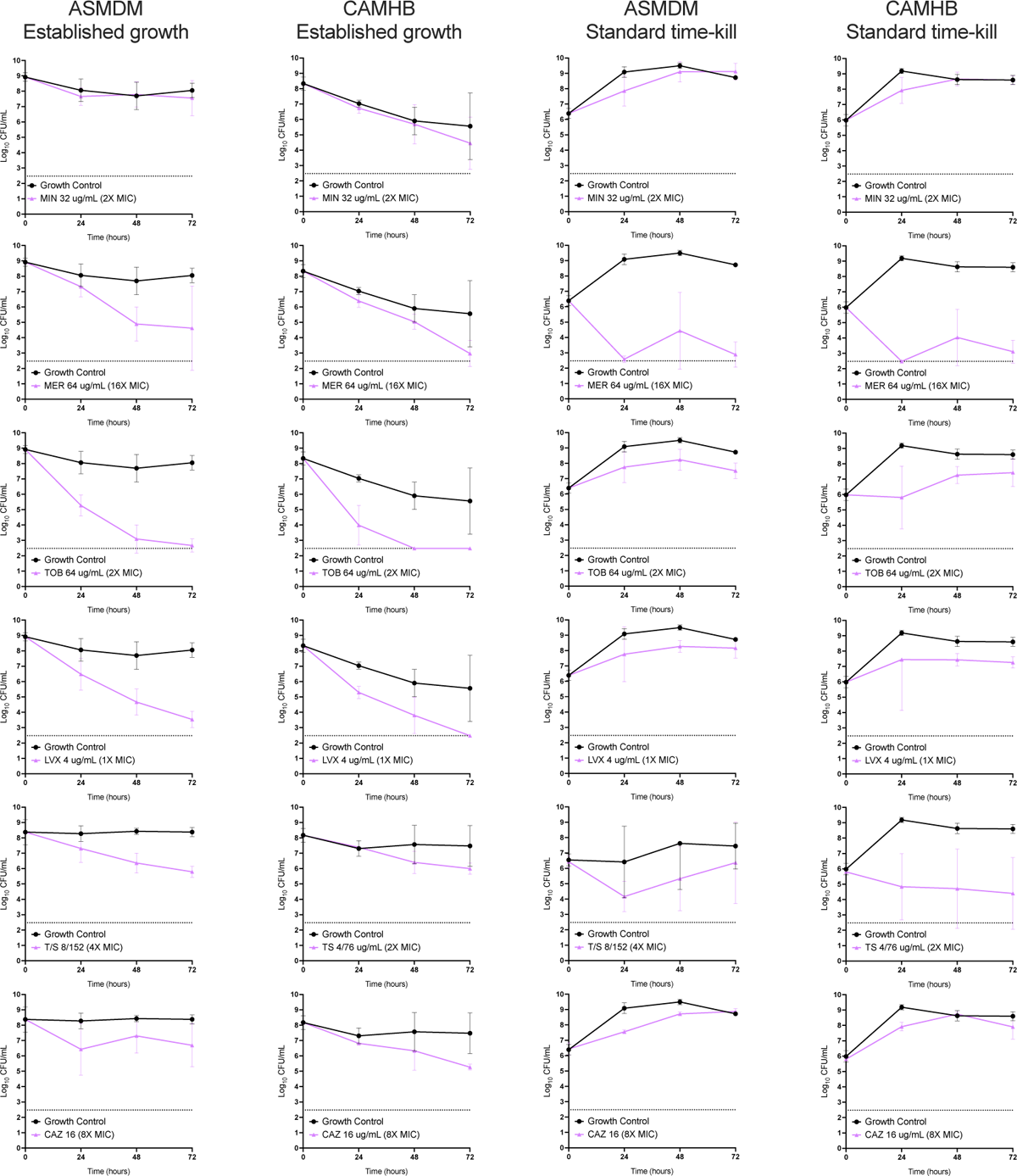
Time-kill studies in different conditions: *B. cenocepacia* K56-2 ASMDM: Artificial sputum medium; CAMHB: cation-adjusted Mueller Hinton broth; MIN: minocycline; MER: meropenem; TOB: tobramycin; LVX: levofloxacin; T/S: trimethoprim-sulfamethoxazole; CAZ: ceftazidime. Graphs show geometric mean and standard deviation of ≥3 replicates. Dashed line indicates assay lower limit of detection. For assays performed simultaneously, the same growth control wells were used.

**FIGURE 3.**
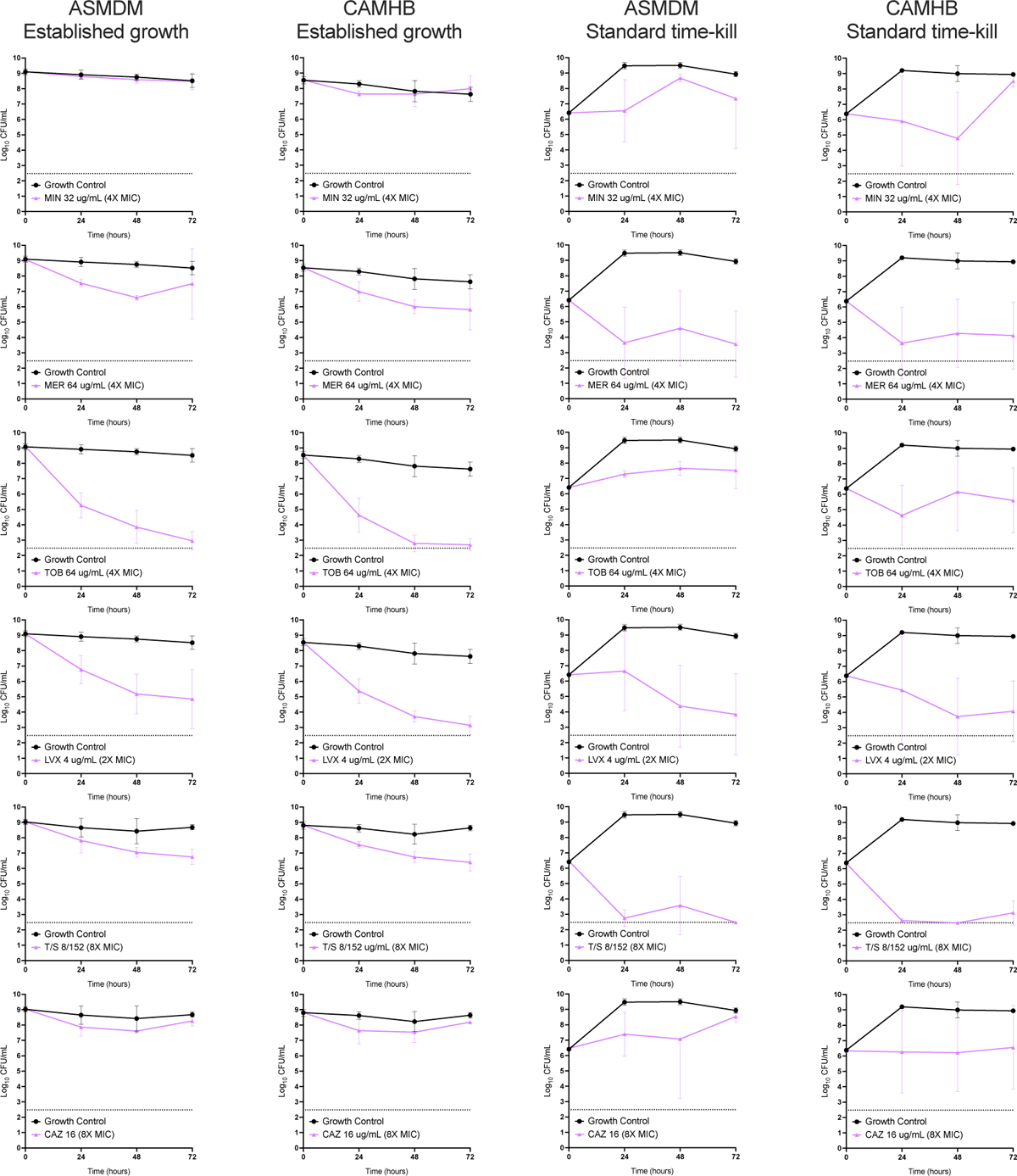
Time-kill studies in different conditions: *B. multivorans* CGD1 ASMDM: Artificial sputum medium; CAMHB: cation-adjusted Mueller Hinton broth; MIN: minocycline; MER: meropenem; TOB: tobramycin; LVX: levofloxacin; T/S: trimethoprim-sulfamethoxazole; CAZ: ceftazidime. Graphs show geometric mean and standard deviation of ≥3 replicates. Dashed line indicates assay lower limit of detection. For assays performed simultaneously, the same growth control wells were used.

### Metabolic activity of bacteria based on microcalorimetry measurements

Bacterial growth in artificial sputum medium is believed to involve establishment of bacterial microcolonies, which have some similarities to biofilms but involve attachment of cells to each other within media rather than to an inanimate surface (21). As an alternate means of observing bacterial response to antibiotics, we used a microcalorimetry instrument to detect bacterial metabolic activity. As can be seen in **Fig. 4**, in which metabolic activity of several different strains was observed immediately following addition of a diluted inoculum (as in standard MIC or time-kill testing), different genera and species have different characteristics “shapes” to their heat profile over time. Both and *B. multivorans* and *B. cenocepacia* reached peak metabolic activity later in ASMDM than in CAMHB, suggesting that the strains are affected by the different media types in a way that could not be appreciated by standard growth curves. **Fig. 5** demonstrates metabolic activity after addition of antibiotics to bacteria that had been growing in ASMDM for 72 hours, as in time-kill curves shown in **Fig. 2**. The elevation in metabolic activity seen over the first 24 hours in growth controls, which likely represents bacterial response to a change in environment during removal of vial caps for addition of antibiotics, is different between the two strains, with *B. multivorans* reaching peak metabolic activity more quickly than *B. cenocepacia* but showing a lower peak value.

**FIGURE 4.**
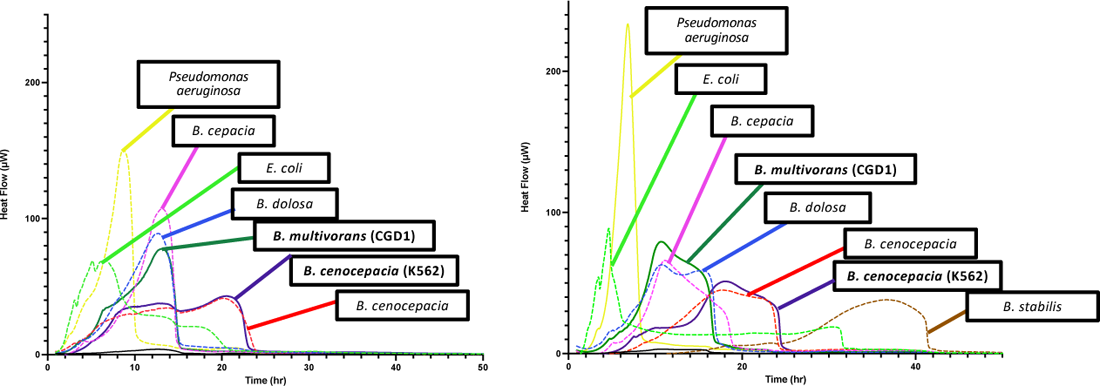
Metabolic activity measured for 50 hours represented as heat flow of bacterial strains grown in ASMDM (L) and CAMHB (R) detected by the CalScreener instrument illustrates differences in growth patterns between media

**FIGURE 5.**
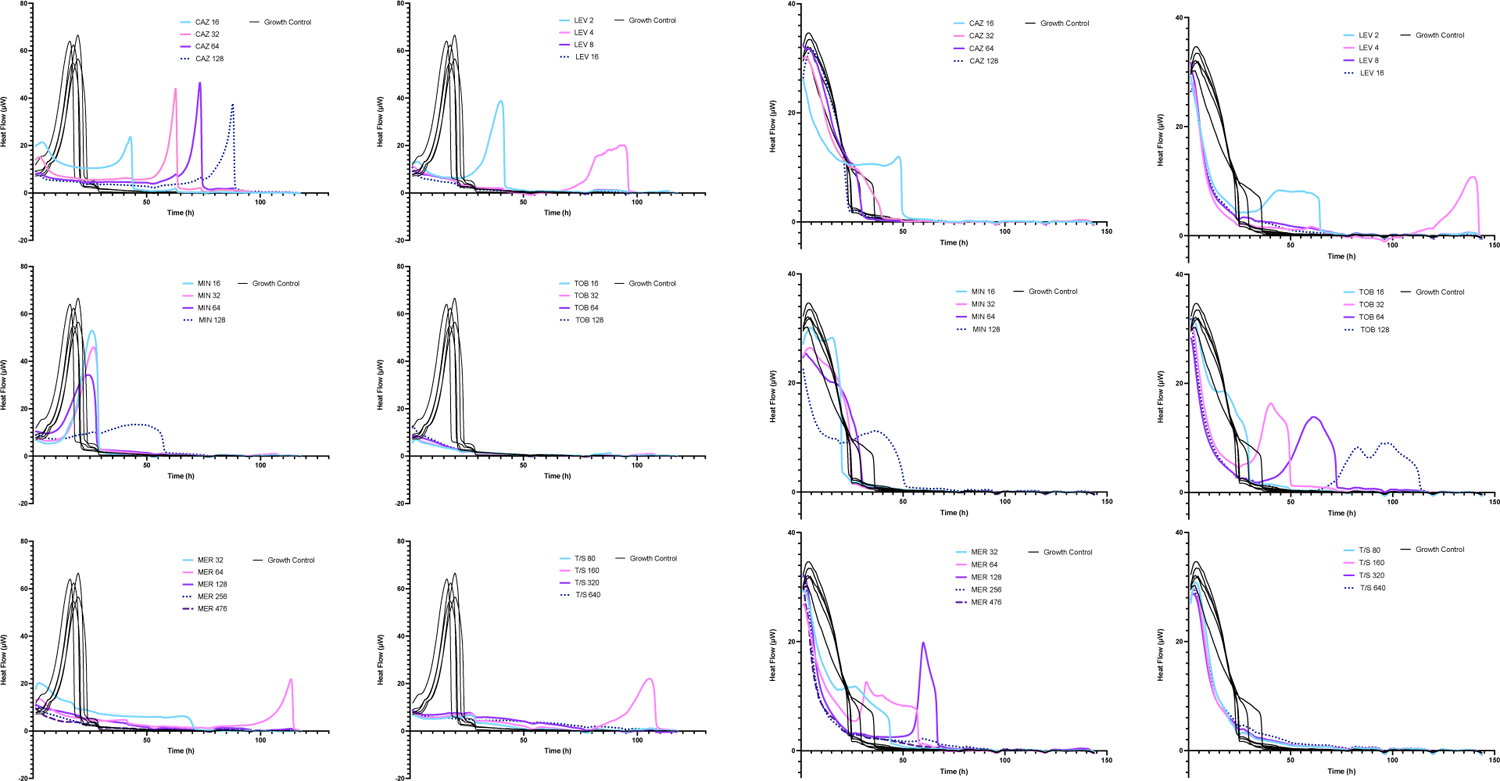
Metabolic activity (heat flow) of bacteria grown in ASMDM for 72 hours prior to addition of antibiotics. (a) *B. cenocepacia* K56-2; (b) *B. multivorans* CGD1. MIN: minocycline; MER: meropenem; TOB: tobramycin; LVX: levofloxacin; T/S: trimethoprim-sulfamethoxazole; CAZ: ceftazidime. Concentrations are in μg/mL. T/S concentrations are expressed as the sum of the two components.

The effects of antibiotics on metabolism in the two strains also differed in ways that were not predictable from time-kill study results. For example, while trimethoprim-sulfamethoxazole exhibited similar activity against the two strains in time-kill studies, with a gradual decline in colony count over 72 hours (**Fig. 2**), the effects of this drug on metabolic activity were markedly different between the two strains, with suppression of the peak in *B. cenocepacia* but only a slightly faster decline in the peak in *B. multivorans*. Similarly, while the effect of tobramycin on the two stains was similar in time-kill studies (**Fig. 2**), the effect of the drug on metabolic activity of *B. cenocepacia* was immediate, with near-complete inhibition of activity by 24 hours even at 0.5X MIC (16 μg/mL), whereas for *B. multivorans*, it had minimal effect until 2X MIC (32 μg/mL); at and above this concentration, it caused a more rapid decline in the initial metabolic peak, followed by later increases in metabolic activity.

### Antibiotic stability

Delayed peaks in metabolic activity microcalorimetry measurements at or beyond 50 hours with some drugs (ceftazidime, and, at lower concentrations, levofloxacin, meropenem and trimethoprim-sulfamethoxazole for *B. cenocepacia*; levofloxacin, meropenem, and tobramycin at some concentrations for *B. multivorans*) (**Fig. 6**) prompted consideration of the possibility of antibiotic breakdown. A biological assay of antibiotic stability assay (**Table S1**) showed that levofloxacin and tobramycin were most stable, with no more than a single two-fold change in concentration at 72 hours, whereas meropenem, ceftazidime, and trimethoprim-sulfamethoxazole showed significant breakdown of >95%.

## DISCUSSION

*Burkholderia cepacia* complex organisms are important pathogens that cause life-threatening disease in people with cystic fibrosis and chronic granulomatous disease, among other conditions. Antibiotic treatment is essential for patients with Bcc infections, yet limitations of standard AST for these organisms have been widely noted, both in terms of their reproducibility and the ability to predict clinical outcomes (14, 15), and evidence-based guidelines for optimal treatment of Bcc do not exist (22). Our results reinforce these concerns. Based on MIC results, a clinician treating a patient infected with the isolates we studied here might well have selected ceftazidime, which has an MIC of 2 μg/mL for both strains, which is well within the CLSI susceptible range of ≤8 μg/mL (20). However, ceftazidime showed limited activity, even at 16X the MIC, in an assay designed to simulate bacterial growth in the lungs of people with CF (**Fig. 2**). Similarly, minocycline had no activity at up to 8X MIC. Although these strains had minocycline MICs in the intermediate and resistant range, such that this drug would not have been a first-line treatment option, the lack of correlation between broth microdilution MIC and activity in the assays employed here raises significant concern regarding the predictive value of MIC testing.

Bcc are considered intrinsically resistant to aminoglycosides because the MICs of these drugs are higher than those safely attainable systemically. However, inhaled tobramycin is routinely used in people with CF as an approach to reduce the pulmonary burden of *Pseudomonas aeruginosa* (23), and when administered in an inhaled formulation, tobramycin reaches alveolar concentrations of up to 1000 μg/mL (24). Our results indicate that in high-inoculum bacterial growth in conditions similar to those found in the CF lung, tobramycin may be particularly effective, suggesting potential therapeutic utility of inhaled tobramycin for people with CF who are colonized with Bcc. Indeed, tobramycin, which is a protein synthesis inhibitor, was more active against established, high-inoculum growth than in standard time-kill experiments, perhaps as a result of metabolic constraints of stationary-phase, nutritionally depleted bacteria. A similar effect was observed with the activity of levofloxacin against the *B. cenocepacia* strain. Interestingly, while antibiotic activity differed in many cases between established growth and standard time-kill assay conditions, antibiotic activity was remarkably similar between media types (ASMDM and CAMHB). This is likely due in part to the fact that, inevitably, ASMDM does not perfectly recapitulate the environment in the CF lung, but also suggests that bacterial density and growth phase may be more important contributors to antibiotic activity than specific micro- and macro-molecular components of the environment.

There are some limitations to the model we used. Nutrients in the media would inevitably have been depleted over the prolonged time course of the experiments, although a nutrient-starved condition may in many ways better replicate conditions in chronic, high-inoculum infections found in the CF lung (25). Furthermore, because this was a single-compartment model, we did not replace antibiotics during the experiment. As a result, drugs that are less stable (ceftazidime, meropenem, trimethoprim-sulfamethoxazole) broke down significantly by the 72 hour time point. Interestingly, however, we did not see significant regrowth in later time periods of most of the experiments even with these drugs, perhaps as a result of a post-antibiotic effect (26). Future experiments utilizing tools such as the hollow-fiber infection model, which allows for removal of waste products, repletion of nutrients, and replacement of antibiotics over time, will allow for characterization of antibiotic activities under conditions that further approach those found *in vivo*.

The European Committee on Antimicrobial Susceptibility Testing (EUCAST) considers the value of MIC testing for Bcc to be so limited that it does not promulgate interpretive criteria for these organisms (13). Our results would seem to support the limitations of standard MIC testing and the need for development of new, more clinically relevant testing methods. One of the primary reasons that organizations may hesitate to issue breakpoints that do not have established clinical predictive value is the possibility that a clinician will believe an antibiotic to be more effective than it is, as appears likely to be the case for ceftazidime in the strains we evaluated here. However, it is worth noting that some drugs, including tobramycin, actually performed better in conditions resembling the CF lung than in standard time-kill studies. More clinically relevant susceptibility testing approaches may, therefore, turn out to open some new treatment doors for patients even as they close others.

## MATERIALS AND METHODS

### Bacterial Isolates

*Burkholderia cenocepacia* isolates K56-2 and *Burkholderia multivorans* isolate CGD1 were obtained from BEI resources (Manassas, VA). *E. coli* ATCC 25922 was obtained from the American Type Culture Collection (Manassas, VA). Strains were colony purified, minimally passaged, and stored at 80°C in tryptic soy broth with 50% glycerol until the time of use.

### Antibiotics and MIC Testing

Stocks of minocycline and ceftazidime (Chem-Impex Intl., Wood Vale, IL), meropenem (GoldBio, St. Louis, MO), tobramycin (Research Products Intl., Mt. Prospect, IL), levofloxacin (Alfa Aesar, Haverhill, MA) and trimethoprim-sulfamethoxazole (RPI, Mount Prospect, IL; Chem-Impex) were prepared according to Clinical and Laboratory Standards Institute (CLSI) guidelines (20), with the following alterations to allow for proper fluid handling by the D300 digital dispenser instrument (HP, Inc., Palo Alto, CA): 0.3% polysorbate 20 (Sigma-Aldrich, St. Louis, MO) was added to aqueous stocks, and trimethoprim and sulfamethoxazole stocks were prepared in DMSO. All antibiotic stock solutions were quality control (QC) tested with *E. coli* ATCC 25922 prior to use. MIC and QC testing were performed in 384-well plates using a digital dispensing-based adaptation of standard CLSI methodology previously developed and validated in our laboratory (27, 28). Antimicrobial solutions were stored as aliquots at 20°C (except ceftazidime, which was stored at −80°C) and were discarded after a single use.

### Preparation of ASMDM

Artificial sputum medium (“ASMDM”) was prepared based on the protocol described by Fung et al. (16)by adding 10 g porcine stomach mucin II (Sigma-Aldrich), fish sperm DNA (Serva, Heidelberg, Germany), 5.9 mg DTPA, 5g NaCl, 2.2 g KCl and 250 mg of each amino acid except tryptophan to 1L of water (pH 6.9). L-tyrosine was dissolved in deionized water and L-cysteine in 0.5 M NaOH before adding. The mixture was autoclaved to sterilize (29). After the mixture had cooled, 10 g of bovine serum albumin (Fisher Bioreagents) and 250 mg of tryptophan, dissolved in water and filter sterilized, were added, along with 5 mL of egg yolk emulsion (16, 17, 21). ASMDM was stored at 4°C until the time of use.

### Time-kill Studies

Bacteria were streaked from frozen stocks onto TSA/5% sheep blood agar plates and incubated overnight at 37°C in ambient air. A starting inoculum of ∼5×10^6^ CFU/mL was prepared in ASMDM or cation-adjusted Mueller-Hinton broth (CAMHB), and 100 μL of this suspension was added to wells in black flat-bottom tissue culture-treated 96-well plates. Plates were incubated at 37°C in ambient air inside a “supermoat” system designed to minimize evaporation (**Fig. S1**). For time-kill studies in established growth, bacteria were incubated for 72 hours prior to addition of antibiotics; for “traditional” time-kill assays, antibiotics were added to wells after the initial time-point was obtained (3 hours after addition of bacteria to wells; see below). Antibiotics were added to wells with the D300 digital dispenser (28).

Bacterial activity was evaluated at 0, 24, 48, and 72 hours after addition of antibiotics by two methods: (1) calculation of percent viability according to capacity of cells to reduce resazurin based on fluorescence measurement and (2) plate count colony enumeration. A separate set of wells was used for each time point. For resazurin-based readings, 10 μL of 100 mg/mL cellulase in 0.05 M citrate buffer (17) was added to wells two hours before the reading time-point to disrupt biofilms and microcolony aggregates, and 0.02% resazurin was added one hour before the time-point. Fluorescence was measured using an excitation wavelength of 540 nm and an emission wavelength of 590 nm. Average background fluorescence from wells containing media alone was subtracted from the value of each well to obtain a normalized reading, and percent viability was determined relative to a well containing bacteria without antibiotic. The contents of each well was then removed for colony enumeration. A 10-fold dilution series was prepared in 0.9% sodium chloride and a 10 μL drop from each dilution was transferred to a Mueller-Hinton plate and incubated overnight in ambient air at 37°C (30). For countable drops (drops containing 3 to 30 colonies), the cell density of the sample was calculated; if more than one dilution for a given sample was countable, the cell density of the two dilutions was averaged. If no drops were countable, the counts for consecutive drops above and below the countable range were averaged. The lower limit of quantitation was 300 CFU/mL.

### Microcalorimetry measurements

The isothermal microcalorimetry instrument SymCel CalScreener^TM^ (Stockholm, Sweden) was used to quantify bacterial metabolism through detection of heat under different conditions (31, 32). Bacteria were streaked from frozen stocks on TSA/5% sheep blood agar plates and incubated overnight at 37°C in ambient air. A starting inoculum of 5×10^6^ CFU/mL was prepared from the overnight plates in ASMDM or CAMHB. Two hundred microliters of this suspension was added to each of 32 plastic inserts in a 48-well CalScreener plate and 10X PBS added to the remaining vials as thermodynamic references. Inserts were then transferred to titanium vials and lids were screwed on according to instrument manufacturer’s instructions. For growth curve studies without antibiotics, microcalorimetry monitoring was initiated at this point and continued for 50 hours. For analysis of antibiotic activity on established bacterial growth, bacteria were added to vials as above, then allowed to grow inside the instrument for 72 hours at 37°C. The plate was then removed from the instrument and lids were removed from vials containing bacteria. Antibiotics were dispensed into the vials using the D300 digital dispenser, lids were replaced, and the plate was put back into the CalScreener for microcalorimetry monitoring for 150 hours, with heat values over time recorded by the instrument.

### Antibiotic Stability Assay

Contents from selected wells from the growth detection and microcalorimetry assays were frozen at −20°C after the completion of the assays for additional experiments. Samples were thawed on the day of the experiment and centrifuged for 2 minutes to pellet bacteria. Supernatant from each tube was filter-sterilized and diluted in CAMHB, based on the predicted drug concentration, to include the QC range for *E.coli* ATCC 25922. Serial two-fold dilutions were performed in CAMHB in an untreated 96-well plate to prepare a standard doubling dilution broth microdilution plate. Antibiotics of interest were also dispensed from frozen stocks using the D300 to prepare a reference plate in parallel. *E. coli* 25922 was added to wells at a final concentration of 5×10^5^ CFU/mL per CLSI guidelines (20) before incubating overnight at 37°C. The MIC of *E. coli* 25922 in the reference plate was used to calculate the starting concentration of antibiotic in each sample in the experimental plate.

---

**FIGURE S1.**
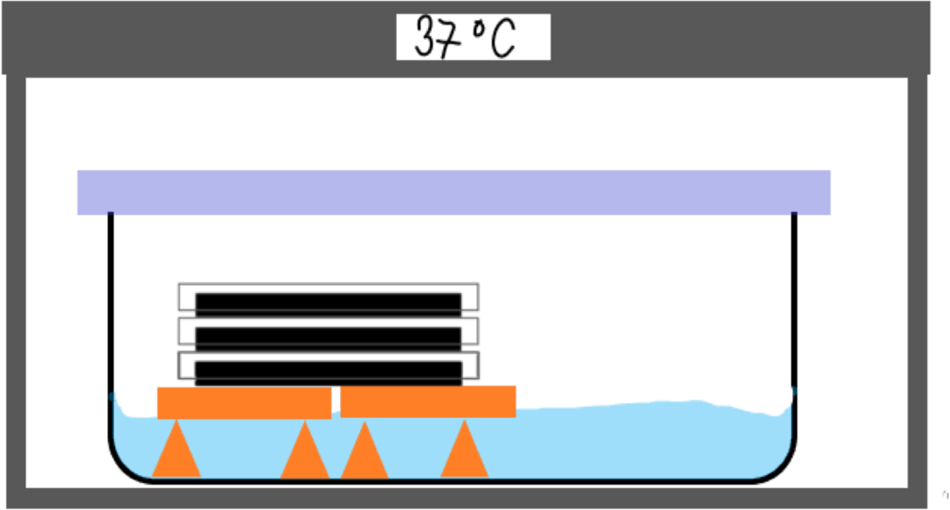
A “supermoat” system for bacterial growth. Bacteria were grown in 96-well plates placed within a container with water at about an inch high to prevent evaporation during multiday experiments. Container was sealed but not airtight.

**FIGURE S2.**
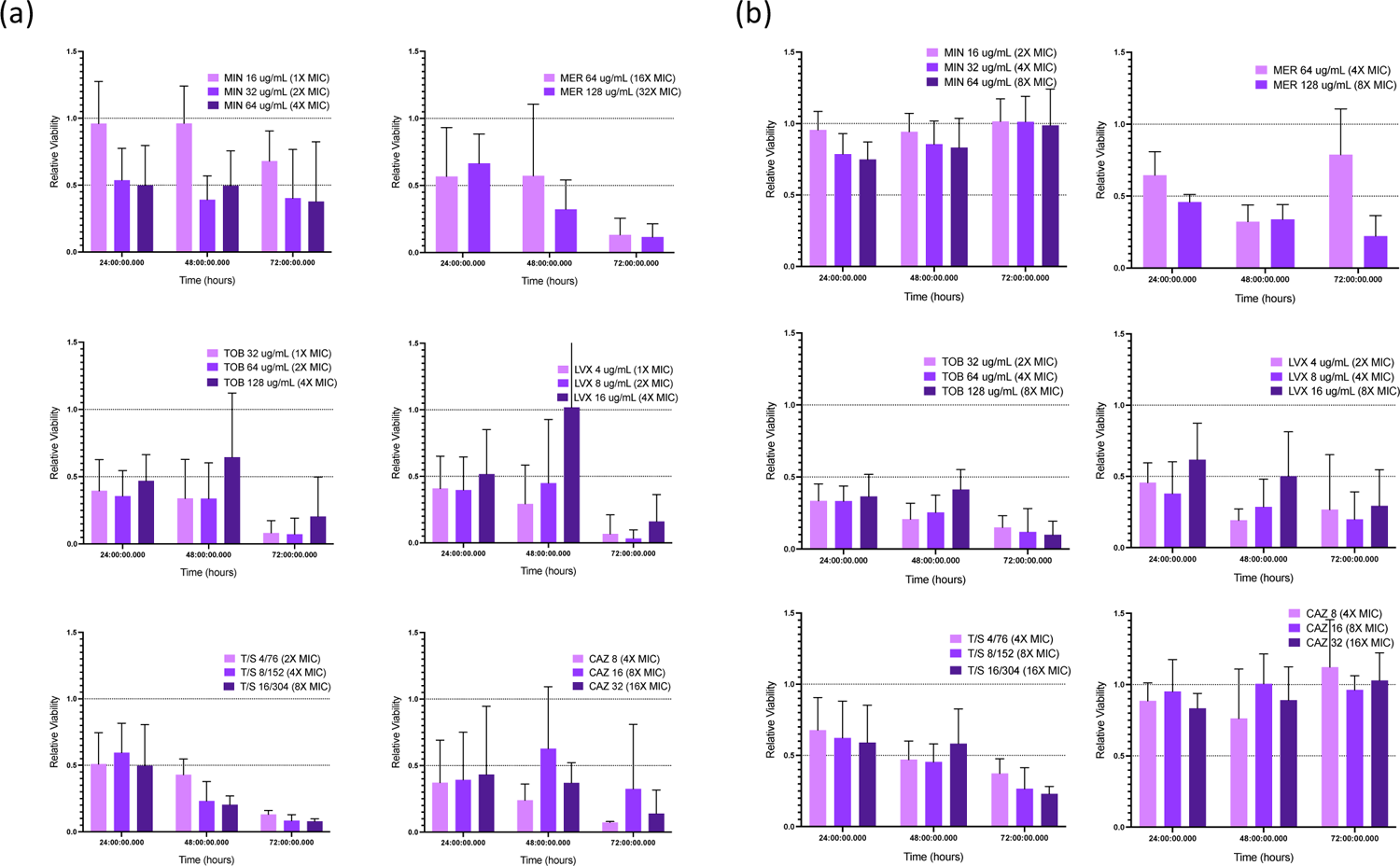
Viability of bacteria in ASMDM with established growth of (a) *B. cenocepacia* K56-2 and (b) *B. multivorans* CGD1 determined by resazurin reduction. MIN: minocycline; MER: meropenem; TOB: tobramycin; LVX: levofloxacin; T/S: trimethoprim-sulfamethoxazole; CAZ: ceftazidime.

**Table S1.** Antibiotic Stability.

| Antibiotic | Strain Treated | Starting Concentration (µg/mL) | Final Concentration (µg/mL) |
| --- | --- | --- | --- |
| Minocycline | *B. cenocepacia* | 64 | 32 |
|  | *B. multivorans* | 64 | 16 |
| Meropenem | *B. cenocepacia* | 128 | 0.5 |
|  | *B. multivorans* | 128 | 0.5 |
| Tobramycin | *B. cenocepacia* | 128 | 64 |
|  | *B. multivorans* | 128 | 64 |
| Ceftazidime | *B. cenocepacia* | 32 | 0.25 |
|  | *B. multivorans* | 32 | 0.5 |
| Levofloxacin | *B. cenocepacia* | 16 | 16 |
|  | *B. multivorans* | 16 | 16 |
| Trimethoprim-Sulfamethoxazole | *B. cenocepacia* | 304/16 | 4.75/0.25 |
|  | *B. multivorans* | 304/16 | 9.5/0.5 |

